# Light-emitting diode-based transcranial photoacoustic measurement of sagittal sinus oxyhemoglobin saturation in hypoxic neonatal piglets

**DOI:** 10.1101/2020.08.22.262451

**Authors:** Jeeun Kang, Raymond C. Koehler, Shawn Adams, Ernest M. Graham, Emad M. Boctor

**Affiliations:** Department of Radiology and Radiological Science, Johns Hopkins University School of Medicine (JHU SOM), Baltimore, MD 21205, USA; Laboratory for Computational Sensing and Robotics, Johns Hopkins University Whiting School of Engineering, Baltimore, MD 21218, USA; Department of Anesthesiology and Critical Care Medicine, Johns Hopkins University School of Medicine, Baltimore, MD 21205, USA; Division of Maternal-Fetal Medicine, Department of Gynecology-Obstetrics, Johns Hopkins University School of Medicine, Baltimore, MD 21205, USA; Neuroscience Intensive Care Nursery Program, Johns Hopkins University School of Medicine, Baltimore, Maryland, MD 21205, USA

## Abstract

We present a light-emitting diode (LED)-based transcranial photoacoustic measurement (LED-trPA) of oxyhemoglobin (HbO_2_) saturation at superior sagittal sinus (SSS) in hypoxic neonatal piglets. The optimal LED imaging wavelengths and frame averaging scheme were determined based on *in vivo* characterization of transcranial sensitivity. Based on the framework (690/850 nm with >20 frame averaging), graded hypoxia was successfully identified in neonatal piglets *in vivo* with less than 10.0 % of root mean squared error (RMSE). This preclinical study suggests the feasibility of a rapid, cost-effective, and safe LED-trPA monitoring of perinatal hypoxia-ischemia and prompt interventions for clinical use.

## Main text

Photoacoustic (PA) imaging is an emerging hybrid modality providing molecular contrast of light absorbance in biological tissue at submillimeter-scale spatial resolution and centimeter-scale penetration depth [1]. Several transcranial PA sensing methods have been presented to quantify electrophysiologic and neurophysiologic dynamics mostly in rodent brain [2–6]. We recently validated the feasibility of transcranial PA measurement of graded oxyhemoglobin (HbO_2_) saturation changes in the superior sagittal sinus (SSS) in a neonatal piglet model simulating a skull thickness and brain development comparable to those of term human neonates [7]. This validation suggests its high translationability for monitoring the incidence of perinatal hypoxic-ischemic encephalopathy (HIE). However, this methodology still conveys concerns remaining on the use of a class-IV high-energy Nd:YAG laser equipped with a tunable optical parametric oscillator (OPO), regarding laser safety, slow scanning speed, bulkiness, and high cost.

There have been several investigations to make PA imaging system safer, faster, more compact, and affordable. A compact approach for brain sensing has been presented using pulsed laser diode (PLD) technology with up to 1–4 mJ per pulse with ~100-nsec pulse duration and 1kHz pulse-repetition frequency (PRF) [8,9]. However, the study only demonstrated a monochrome or single-point reading, lacking spectral and spatial specificity, respectively. On the other hand, light-emitting diode (LED)-based PA imaging has been receiving attention at the forefront of translational investigations [10]. State-of-the-art LED technology provides discrete near-infrared wavelengths (690, 750, 820, 850, 940, and 980 nm) at up to 200 μJ per pulse with 30–100-nsec pulse duration and 0.2–16KHz PRF. Notably, the LED source offers no need for safety goggles and barriers which are strictly obliged when using high-energy lasers. Furthermore, its unit cost is below 10 % from what is needed for an Nd:YAG OPO laser ($10–15K vs. $70–200K). However, to the best of our knowledge, transcranial sensing applications with LEDs have remained unexplored due to the challenges with high attenuation in the scalp and skull layers. In this short communication, we validate the feasibility of multi-spectral LED-based transcranial PA measurement (LED-trPA) of sagittal sinus HbO2 saturation in neonatal piglets with graded hypoxia *in vivo*. Our LED-trPA imaging system was built with 4-channel pulsed LED source (Prexion Inc., Japan) and ultrasound research package (Vantage 256, Verasonics Inc., USA) (Fig. 1).

**Fig 1.**
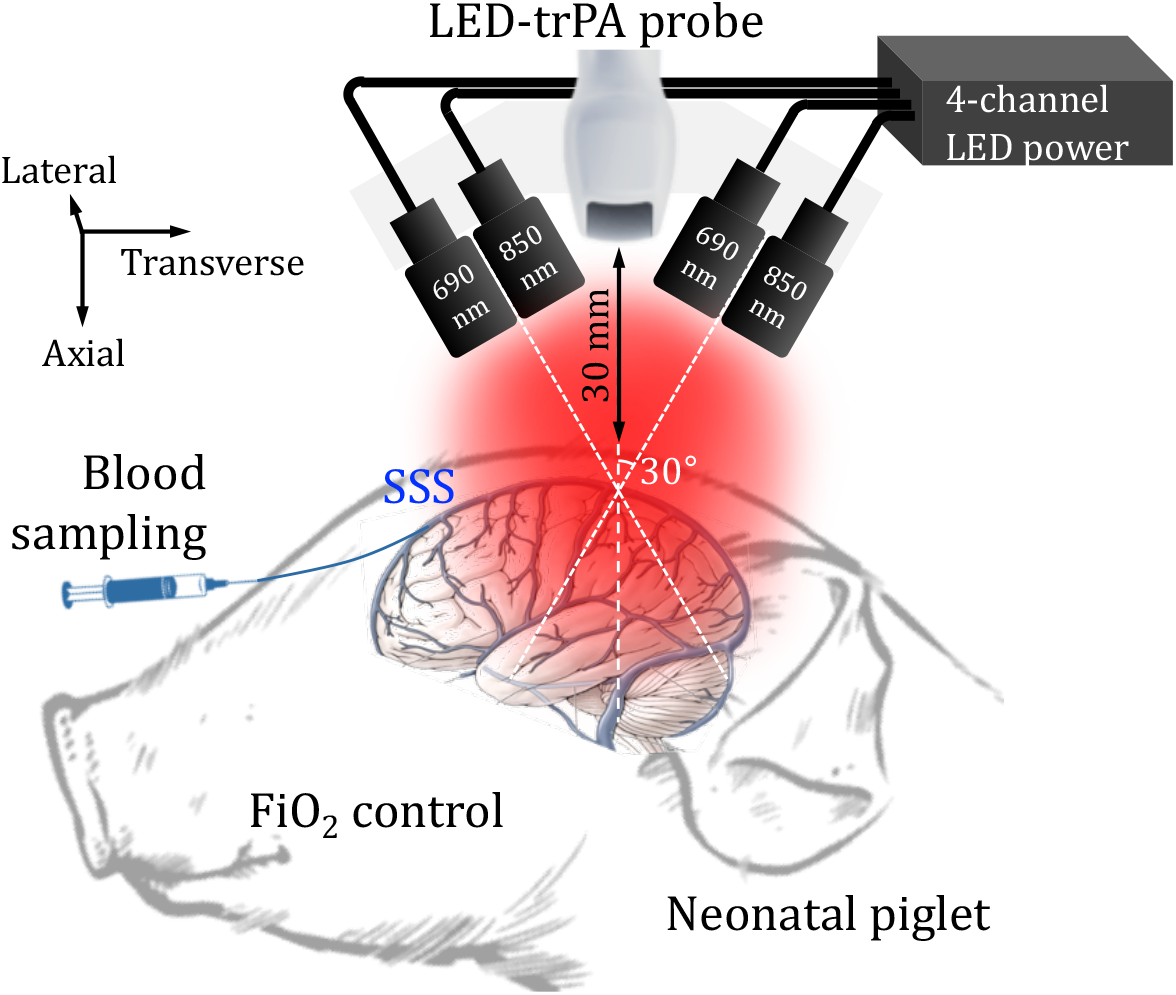
Pulsed LED-based transcranial photoacoustic sensing (LED-trPA) system. Abbreviations: superior sagittal sinus, SSS; fraction of inspired O_2_, FiC_2_.

All procedures on piglets were approved by the Johns Hopkins University Animal Care and Use Committee. Transcranial energy density threshold for SSS sensing 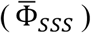 was first characterized in anesthetized and mechanically ventilated 3-5-day-old neonatal piglets (1.5-2.1 kg; fraction of inspired O_2_ (FiO_2_) = 0.33; *n* = 4). In this signal characterization stage, we employed the experimental setup based on an Nd:YAG OPO laser (Phocus Inline, Opotek Inc., USA) to investigate a wide range of incident energy density [7]. Catheters were inserted into the SSS to obtain cortical venous blood samples for blood gas measurement of the ground-truth HbO2 saturation at SSS. The pulsed laser was delivered by an optical fiber bundle attached to a 5.2-MHz linear array (L7-5, ATL Inc., USA). 10 % of the OPO output was intercepted for simultaneous recording. The piglet head was scanned in near-infrared LED wavelengths (i.e., 690, 750, 820, and 850 nm) with graded flash-to-Q-switch delays in the Nd:YAG laser: 200 (optimal), 205, 210, 215 μsec. A flash-to-Q-switch delay longer than 200 μsec lowers the energy density at the fiber output: −16.13, −32.34, and −47.78 % for 205, 210, and 215 μsec in average over entire wavelength range. The 1.5×1.5-mm^2^ region-of-interest (ROI) was selected on SSS to quantify a regional PA spectrum (Fig. 2a). Fig. 2b shows the PA sensitivity at SSS as a function of incident energy density at each wavelength, fitted to linear regression curves (Mathworks Inc., USA). A constraint was applied to intercept the origin point (i.e., no PA intensity at 0 mJ/cm^2^). Note that the background noise level (BG) was measured without laser excitation, and then subtracted from the measured PA intensity. In the examples shown in Fig. 2b with no frame averaging, the linear regression slopes normalized to the BG were 8.06, 7.10, 5.76, and 7.59 (i.e., PA intensity at SSS / mJ · cm^−2^ / BG) at 690, 750, 820, and 850 nm, respectively. This implies that 690 nm and 850 nm yield the higher transcranial sensitivity at the SSS per an energy density than others. The BG was lowered with a factor of 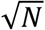, reflecting reciprocal signal-to-noise ratio (SNR) improvement [11,12]. The cross point between the fitted curves to the BG was then identified at each frame averaging scheme (1, 2, 5, 10, 20, 50, 100, 200, 500, 1,000, 2,000 frames), specifying the energy density delivering 0-dB SNR.

**Fig 2.**
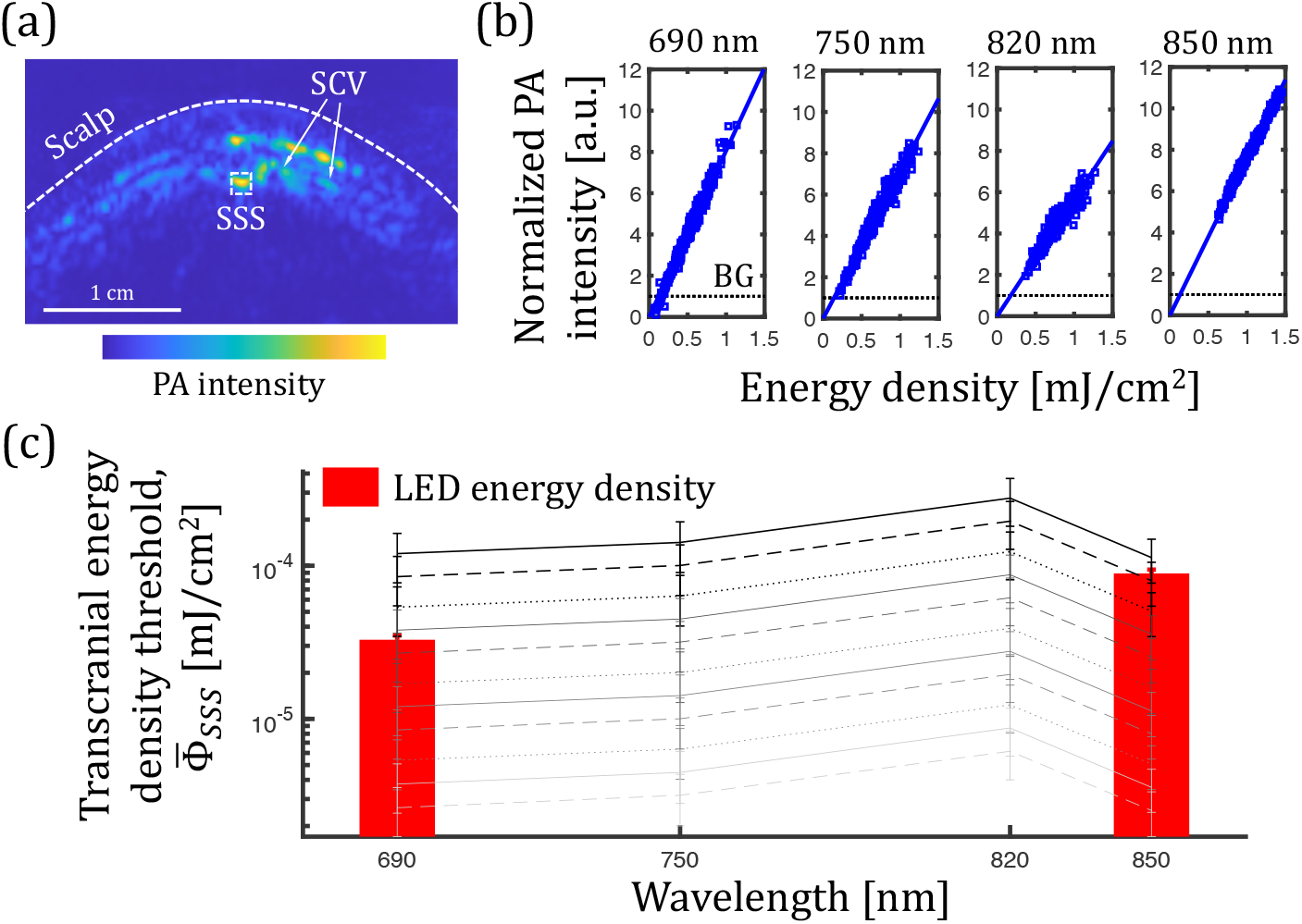
Transcranial energy density threshold for SSS sensing, 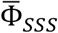 (a) exemplary PA cross-section of piglet head and region-of-interest at SSS. (b) Linear regression curves between energy density and PA intensity at SSS. (c) 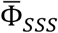 curves defining 0-dB SNR at SSS for 1, 2, 5, 10, 20, 50, 100, 200, 500, 1,000, and 2,000 frame averaging schemes (black to gray). Red bars indicate LED energy densities at 690 and 850 nm. Abbreviations: superior cortical veins, SCV.

Fig. 2c shows the 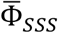 estimated under different frame averaging schemes. The influential factor should be combinational among (1) total hemoglobin absorbance at SSS; (2) optical / acoustic attenuation in scalp and skull layers; and (3) noise level in ultrasound system. In general, the spectral trend of 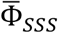 followed the inversion of the total hemoglobin absorbance at the ground-truth HbO_2_ saturation in the piglets, i.e., 52.33 ± 5.95 % (i.e., 0.89, 1.06, 1.23, and 1.13 when ×1,000 at 690, 750, 820, and 850 nm [13]), implying that more hemoglobin absorbance lowers 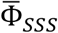. Also, the optical transmission through skull and scalp layers is another significant factor. The skull will have more absorptive attenuation as one gets close to 650 nm and 900 nm, while holding a relatively stable level between 700 nm to 850 nm. [14,15]. On the other hand, the dominant absorber in the scalp layer is melanin which gradually increase optical attenuation as one gets close to shorter wavelength range [16]. The inversely proportional relationship was identified between 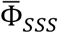 and the number of frames averaged, as expected.

From the 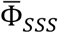 characterization, we selected 690 nm and 850 nm for LED-trPA to secure low 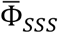 and spectral independency at the same time. When 2 LED arrays were used for each wavelength, the LED-trPA probe yielded the energy density at the depth of subject surface (Φ*_LED_*): 32.86 ± 2.23 and 89.03 ± 5.28 μJ/cm^2^ at 690 nm and 850 nm both with 105 nsec pulse duration (red bars in Fig. 2c). Statistical significance of Φ_LED_ (< 0.05) was obtained from 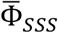 when applied 20- and 5-frame averaging at 690 nm and 850 nm, respectively: (690 nm) 119.99±42.65, 84.85±30.12, 53.67±19.08, 37.98±13.48, 26.87±9.53* (*p*=0.0094), 16.99±6.02, 12.02±4.25, 8.48±3.02, 5.38±1.93, 3.76±1.33, and 2.63±0.93 μJ/cm^2^; (850 nm) 113.00±35.98, 79.90±25.44, 50.55±16.11* (*p*=0.001), 35.73±11.36, 25.28±8.04, 15.98±5.12, 11.28±3.61, 8.00±2.54, 5.08±1.61, 3.58±1.12, and 2.53±0.82 μJ/cm^2^ for 1, 2, 5, 10, 20, 50, 100, 200, 500, 1,000, and 2,000 frame averaging. From the Φ_*LED*_ and 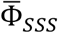, an expected SNR at SSS (i.e., SNR_est_) was calculated at each wavelength and frame averaging scheme, assuming that the PA intensity is proportional to the effective energy density:

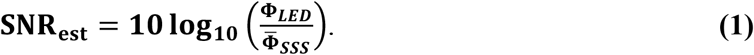

The LED-trPA system configuration was then applied in our following *in vivo* validation study (FiO_2_ = 0.33, *n* = 3). Four LED bars were aligned to a linear array transducer to obtain a coronal cross-section of piglet head (Fig. 1). Fig. 3a shows the LED-trPA images at 690 and 850 nm with 1, 20, 200, 2,000 frame averaging schemes. The 1.5×1.5- mm^2^ SSS region was again selected for the quantification of SNR:

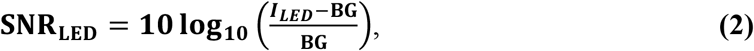

where *l*_LED_ is the LED light-induced PA intensity at SSS. Fig. 3b and Table 2 presents the SNR_est_ and SNRLED together for each frame averaging scheme at 690 nm and 850 nm, respectively. Overall, the LED-trPA measurements showed a well-matched trend to the corresponding estimations over different frame averaging schemes with 0.35±0.43 dB and 1.06±0.12 dB of differences (mean ± standard deviation, SD) at 690 nm and 850 nm, respectively.

**Fig 3.**
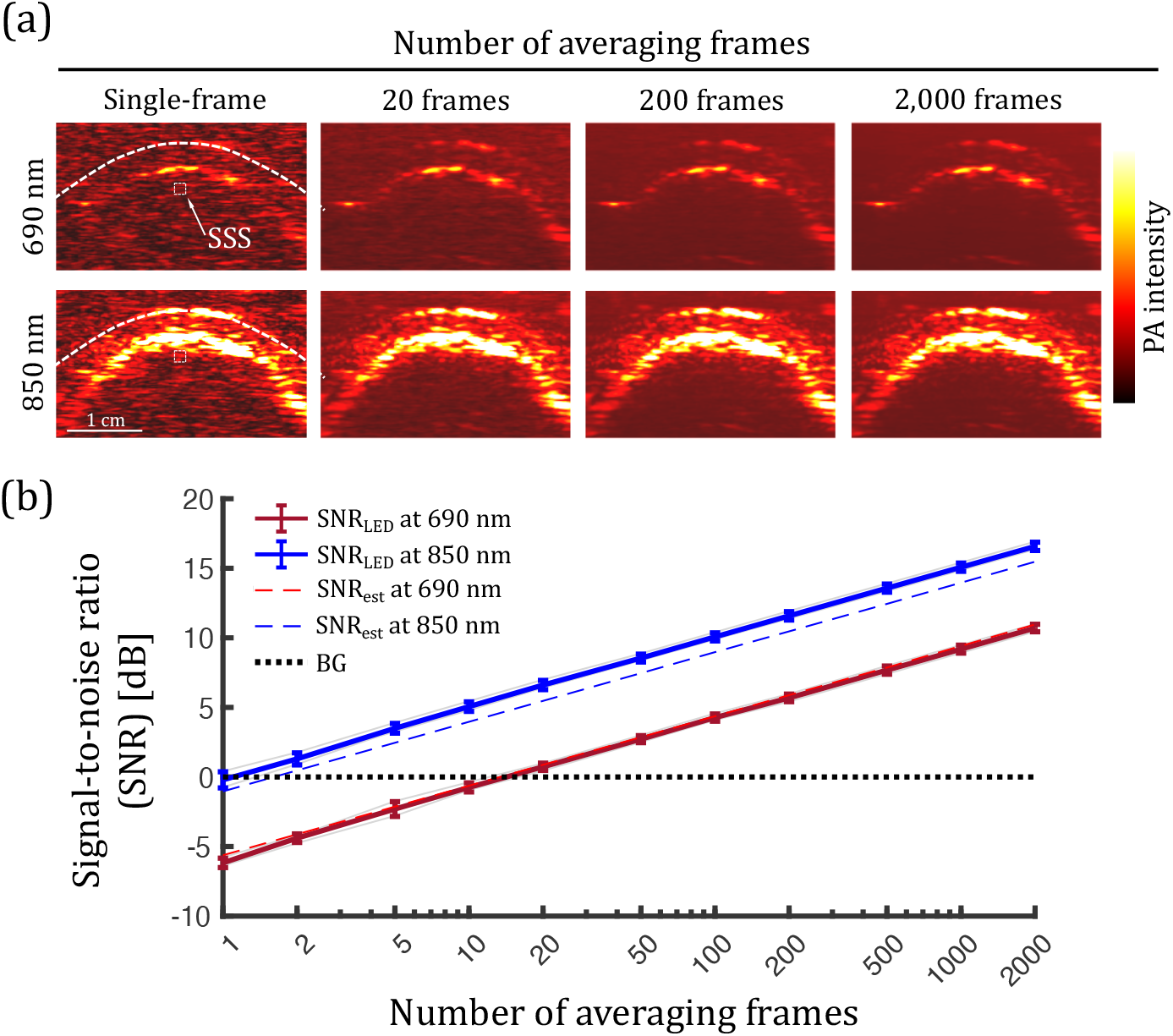
(a) LED-trPA images at 690 nm and 850 nm, processed with 1, 20, 200, 2,000 frame averaging schemes, (b) Comparison between SNR_est_ and SNR_LED_ at 690 nm and 850 nm.

We then induced graded hypoxia in the neonatal piglets by progressively changing FiO_2_ level: 1.00, 0.80, 0.60, 0.50, 0.40 0.33, 0.20, 0.18, 0.16, 0.15. At each FiO_2_ step, a 5-min stabilization period was allowed before the LED-trPA. 2,000 LED-trPA imaging frames were obtained each, followed by the collection of arterial and SSS blood samples before translating to the next FiO2 level. In some experiments, data were not obtained due to low MAP or inability to draw blood from the SSS catheter. From the data collected, linear regression curves were reconstructed between ground truths (blood gas measurements) and the LED-trPA-based HbO_2_ saturation estimations (Fig. 4a). With 2,000-frame averaging, the linear regression slope was 0.77 and y-intercept at 10.54 % with the goodness of fit (R^2^) at 0.80. Root mean squared error (RMSE) to the ground truth (unity curve) was 6.22 %, and the Bland-Altman estimation analysis presented no physiologically meaningful bias, having −1.38±12.15 % (mean±1.96 SD). The deviation of the accuracy was repeatedly evaluated for 1, 2, 5, 10, 20, 50, 100, 200, 500, 1,000 frame averaging schemes with 20 permutations. Fig. 4c shows the changes in slope, y-intercept, R^2^, and RMSE for the linear regression curves with different frame averaging schemes. In general, more frame averaging yielded the progressively improved HbO_2_ saturation estimation. Notably, single-frame LED-trPA imaging failed to track the change in HbO_2_ saturation at SSS with 0.38 ± 0.18 of slope, 31.34 ± 8.14 of y-intercept, 0.19 ± 0.13 of R^2^, and 15.61 ± 2.10 of RMSE. Otherwise, the 20-frame averaging scheme, enabled the first positive SNR at SSS for both wavelengths (Fig. 3b), derived the first estimation point limiting the RMSE below 10 % in HbO_2_ saturation (i.e., 8.44 ± 1.03, mean ± SD). However, the linear regression curve still presented insufficient accuracy to track HbO_2_ saturation changes at SSS: 0.67 ± 0.06 of slope, 12.49 ± 2.21 of y-intercept, 0.62 ± 0.09 of R^2^. This can be explained by still insufficient SNR at SSS (i.e., 0.87 dB) with 690-nm LED. From 50- to 1,000-frame averaging scheme, the gradual increase of correlation accuracy to ground truth was found, agreed with Bland-Altman estimation analysis showing no physiologically meaningful bias observed at low or high HbO_2_ saturation levels.

**Fig 4.**
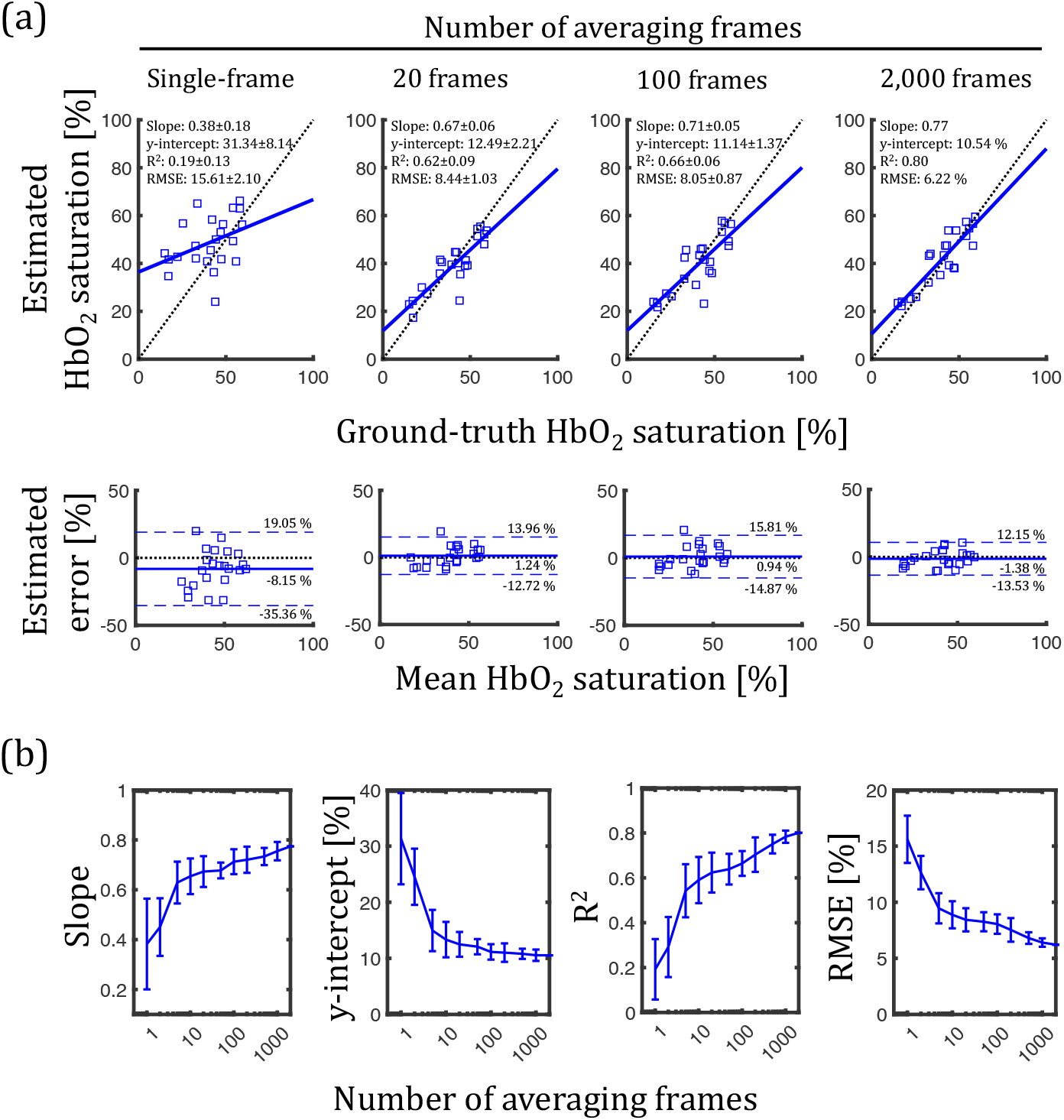
Correlation of estimated and ground-truth HbO_2_ saturation at SSS in neonatal piglets. (a) Linear regression curves and Bland-Altman estimation analysis for different frame averaging schemes. (b) Slope, y-intercept, R^2^ and RMSE of the linear regression curves.

In this study, we presented framework for a non-invasive, real-time, and safe LED-trPA of SSS in neonatal piglets *in vivo* as a significant translational step towards a rapid detection of perinatal HIE, enabling immediate interventions for tissue salvage. We first characterized 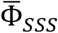the minimal energy density threshold for SSS sensing, 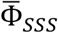, followed by *in vivo* LED-trPA evaluation of graded hypoxia in the neonatal piglets. The use of the neonatal piglet model provides skull thickness and brain development comparable to those in term human neonates [17,18]. Moreover, the pig has skin more similar to humans than rodents [19].

Several aspects of this approach will need to be considered for improving this methodology: (1) Even though the LED-trPA framework could provide RMSE less than 10%, the linear regression presented limited accuracy when compared to our previous results with Nd:YAG OPO laser, even with 2,000 frame averaging: 0.77 vs. 1.00 in slope; 10.54 % vs. 1.49 % in y-intercept; 0.80 vs. 0.87 of R^2^ [7]. It may be suggesting more frame averaging, but we concern the error in spectral decomposition given the much broader linewidth in the LEDs than that in the Nd:YAG OPO laser (20-35 nm vs. 2-3 nm). It could be also the source of error in our SNR estimation (Fig. 3). The appropriate correction in spectral decomposition will enhance the estimation accuracy. (2) The abundant temporal information in LED-trPA opens another opportunity for further enhancement. For instance, our group presented a deep neural network that improved the LED-based PA imaging quality by employing features in temporal sequences [20]. The solution may significantly improve the sensitivity of LED-trPA. However, a great caution will be needed to preserve the spectral features at individual wavelengths. (3) Embedding the LED-trPA on ultra-compact ultrasound systems will facilitate its clinical translation [21,22]. In this integrative phase, strict considerations on thermal and electrical interferences would be needed to be in a compact form factor.

**Table 1.** Expected and measured signal-to-noise ratio (SNR) at SSS *in vivo* for different frame averaging schemes. The asterisk marks follow the earliest point with statistical significance between 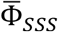 and Φ_LED_.

| | $N$ | 1 | 2 | 5 | 10 | 20 | 50 | 100 | 200 | 500 | 1,000 | 2,000 |
| --- | --- | --- | --- | --- | --- | --- | --- | --- | --- | --- | --- | --- |
| 690 nm | $SNR_{est}$ | -5.62 | -4.12 | -2.13 | -0.63 | 0.87* | 2.86 | 4.37 | 5.88 | 7.86 | 9.42 | 10.96 |
| | $SNR_{LED}$ | -6.17 ± 0.34 | -4.40 ± 0.31 | -2.30 ± 0.53 | -0.75 ± 0.33 | 0.73 ± 0.26* | 2.72 ± 0.25 | 4.27 ± 0.26 | 5.68 ± 0.27 | 7.68 ± 0.29 | 9.19 ± 0.29 | 10.71 ± 0.27 |
| 850 nm | $SNR_{est}$ | -1.03 | 0.46 | 2.45* | 3.96 | 5.46 | 7.46 | 8.97 | 10.46 | 12.44 | 13.96 | 15.47 |
| | $SNR_{LED}$ | -0.21 ± 0.58 | 1.30 ± 0.43 | 3.50 ± 0.35* | 5.06 ± 0.34 | 6.61 ± 0.33 | 8.55 ± 0.28 | 10.07 ± 0.29 | 11.58 ± 0.30 | 13.57 ± 0.29 | 15.08 ± 0.30 | 16.58 ± 0.30 |

